## Supplementary Table 1 for "Genome assembly of the numbat (*Myrmecobius fasciatus*), the only termitivorous marsupial"

**Supplementary Table 1.** Accession numbers for sequences used as BLAST queries and to generate phylogenetic trees in Figure 3, 5 and 6

| Gene family | Gene ID | Accession number | Corresponding figure |
|---|---|---|---|
| TS1R | PhciTAS1R1 | GCA_002099425.1 | Figure 3 |
| TS1R | SahaTAS1R1 | GCA_000189315.1 | Figure 3 |
| TS1R | ModoTAS1R1-partial | GCA_000002295.1 | Figure 3 |
| TS1R | HosaTAS1R1 | Q7RTX1 | Figure 3 |
| TS1R | MumuTAS1R2 | Q925I4 | Figure 3 |
| TS1R | HosaTAS1R2 | Q8TE23 | Figure 3 |
| TS1R | PhciTAS1R2 | GCA_002099425.1 | Figure 3 |
| TS1R | SahaTAS1R2 | GCA_000189315.1 | Figure 3 |
| TS1R | ModoTAS1R2 | GCA_000002295.1 | Figure 3 |
| TS1R | HosaTAS1R3 | Q7RTX0 | Figure 3 |
| TS1R | MumuTAS1R3 | Q925D8 | Figure 3 |
| TS1R | PhciTAS1R3 | GCA_002099425.1 | Figure 3 |
| TS1R | SahaTAS1R3 | GCA_000189315.1 | Figure 3 |
| TS1R | ModoTAS1R3 | GCA_000002295.1 | Figure 3 |
| TS2R | tacAcuTAS2R814_ta | GCA_015852505.1 | Figure 5 |
| TS2R | tacAcuTAS2R813_ta | GCA_015852505.1 | Figure 5 |
| TS2R | tacAcuTAS2R802_ta | GCA_015852505.1 | Figure 5 |
| TS2R | ornAnaTAS2R801_oa | GCF_000002275.2 | Figure 5 |
| TS2R | ornAnaTAS2R813B_oa | GCF_000002275.2 | Figure 5 |
| TS2R | ornAnaTAS2R812_oa | GCF_000002275.2 | Figure 5 |
| TS2R | ornAnaTAS2R813A_oa | GCF_000002275.2 | Figure 5 |
| TS2R | ornAnaTAS2R811_oa | GCF_000002275.2 | Figure 5 |
| TS2R | ornAnaTAS2R810_oa | GCF_000002275.2 | Figure 5 |
| TS2R | ornAnaTAS2R802_oa | GCF_000002275.2 | Figure 5 |
| TS2R | monDomTAS2R1_md | GCA_000002295.1 | Figure 5 |
| TS2R | monDomTAS2R2_md | GCA_000002295.1 | Figure 5 |
| TS2R | monDomTAS2R4A_md | GCA_000002295.1 | Figure 5 |
| TS2R | monDomTAS2R38_md | GCA_000002295.1 | Figure 5 |
| TS2R | monDomTAS2R41_md | GCA_000002295.1 | Figure 5 |
| TS2R | monDomTAS2R60_md | GCA_000002295.1 | Figure 5 |
| TS2R | monDomTAS2R702_md | GCA_000002295.1 | Figure 5 |
| TS2R | monDomTAS2R703_md | GCA_000002295.1 | Figure 5 |
| TS2R | monDomTAS2R704A_md | GCA_000002295.1 | Figure 5 |
| TS2R | monDomTAS2R704B_md | GCA_000002295.1 | Figure 5 |
| TS2R | monDomTAS2R704C_md | GCA_000002295.1 | Figure 5 |
| TS2R | monDomTAS2R705_md | GCA_000002295.1 | Figure 5 |
| TS2R | monDomTAS2R710_md | GCA_000002295.1 | Figure 5 |
| TS2R | monDomTAS2R711A_md | GCA_000002295.1 | Figure 5 |
| TS2R | monDomTAS2R711B_md | GCA_000002295.1 | Figure 5 |
| TS2R | monDomTAS2R711C_md | GCA_000002295.1 | Figure 5 |
| TS2R | monDomTAS2R711D_md | GCA_000002295.1 | Figure 5 |
| TS2R | monDomTAS2R712_md | GCA_000002295.1 | Figure 5 |
| TS2R | monDomTAS2R720A_md | GCA_000002295.1 | Figure 5 |
| TS2R | monDomTAS2R720B_md | GCA_000002295.1 | Figure 5 |
| TS2R | monDomTAS2R722_md | GCA_000002295.1 | Figure 5 |
| TS2R | monDomTAS2R723_md | GCA_000002295.1 | Figure 5 |
| TS2R | monDomTAS2R726A_md | GCA_000002295.1 | Figure 5 |

| | | | |
|---|---|---|---|
| TS2R | monDomTAS2R726B_md | GCA_000002295.1 | Figure 5 |
| TS2R | monDomTAS2R727_md | GCA_000002295.1 | Figure 5 |
| TS2R | monDomTAS2R728_md | GCA_000002295.1 | Figure 5 |
| TS2R | monDomTAS2R729_md | GCA_000002295.1 | Figure 5 |
| TS2R | MumuTAS2R105_mm | Q9JKT4 | Figure 5 |
| TS2R | MumuTAS2R140_mm | Q7TQA4 | Figure 5 |
| TS2R | MumuTAS2R4_mm | Q9JKT3 | Figure 5 |
| TS2R | MumuTAS2R7_mm | P59530 | Figure 5 |
| TS2R | MumuTAS2R40_mm | Q7TQA4 | Figure 5 |
| TS2R | MumuTAS2R143_mm | Q7TQB9 | Figure 5 |
| TS2R | MumuTAS2R136_mm | Q7TQA8 | Figure 5 |
| TS2R | MumuTAS2R134_mm | Q7TQB0 | Figure 5 |
| TS2R | MumuTAS2R119_mm | Q9JKT2 | Figure 5 |
| TS2R | MumuTAS2R38_mm | Q7TQA6 | Figure 5 |
| TS2R | MumuTAS2R16_mm | P59529 | Figure 5 |
| TS2R | MumuTAS2R3_mm | Q7TQA7 | Figure 5 |
| TS2R | MumuTAS2R41_mm | P59532 | Figure 5 |
| TS2R | MumuTAS2R135_mm | Q7TQA9 | Figure 5 |
| TS2R | MumuTAS2R13_mm | Q7TQA4 | Figure 5 |
| TS2R | HosaTAS2R1_hs | Q9NYW7 | Figure 5 |
| TS2R | HosaTAS2R3_hs | Q9NYW6 | Figure 5 |
| TS2R | HosaTAS2R4_hs | Q9NYW5 | Figure 5 |
| TS2R | HosaTAS2R5_hs | Q9NYW4 | Figure 5 |
| TS2R | HosaTAS2R7_hs | Q9NYW3 | Figure 5 |
| TS2R | HosaTAS2R8_hs | Q9NYW2 | Figure 5 |
| TS2R | HosaTAS2R9_hs | Q9NYW1 | Figure 5 |
| TS2R | HosaTAS2R10_hs | Q9NYW0 | Figure 5 |
| TS2R | HosaTAS2R13_hs | Q9NYV9 | Figure 5 |
| TS2R | HosaTAS2R14_hs | Q9NYV8 | Figure 5 |
| TS2R | HosaTAS2R16_hs | Q9NYV7 | Figure 5 |
| TS2R | HosaTAS2R19_hs | P59542 | Figure 5 |
| TS2R | HosaTAS2R20_hs | P59543 | Figure 5 |
| TS2R | HosaTAS2R30_hs | P59541 | Figure 5 |
| TS2R | HosaTAS2R31_hs | P59538 | Figure 5 |
| TS2R | HosaTAS2R38_hs | P59533 | Figure 5 |
| TS2R | HosaTAS2R39_hs | P59534 | Figure 5 |
| TS2R | HosaTAS2R40_hs | P59535 | Figure 5 |
| TS2R | HosaTAS2R41_hs | P59536 | Figure 5 |
| TS2R | HosaTAS2R42_hs | Q7RTR8 | Figure 5 |
| TS2R | HosaTAS2R43_hs | P59537 | Figure 5 |
| TS2R | HosaTAS2R46_hs | P59540 | Figure 5 |
| TS2R | HosaTAS2R50_hs | P59544 | Figure 5 |
| TS2R | HosaTAS2R60_hs | P59551 | Figure 5 |
| TS2R | HosaTAS2R2_hs | Q50KZ9 | Figure 5 |
| TS2R | HosaTAS2R45_hs | P59539 | Figure 5 |
| TS2R | SahaTAS2R7_sh | GCA_000189315.1 | Figure 5 |
| TS2R | SahaTAS2R3-like_sh | GCA_000189315.1 | Figure 5 |
| TS2R | SahaTAS2R38_sh | GCA_000189315.1 | Figure 5 |
| TS2R | SahaTAS2R4_sh | GCA_000189315.1 | Figure 5 |
| TS2R | SahaTAS2R7-like_sh | GCA_000189315.1 | Figure 5 |

| | | | |
|---|---|---|---|
| TS2R | TrvuTAS2R3-like_tv | | Figure 5 |
| TS2R | TrvuTAS2R41-like_tv | | Figure 5 |
| Aquaporin | Koala-AQP1 | A0A6P5ID11 | N/A |
| Aquaporin | Human-AQP1 | P29972 | N/A |
| Aquaporin | Mouse-AQP1 | Q02013 | N/A |
| Aquaporin | Opossum-AQP1 | XM_007505813.1 | N/A |
| Aquaporin | Devil-AQP1 | A0A7N4NY14 | N/A |
| Aquaporin | Koala-AQP2 | A0A6P5IKS2 | N/A |
| Aquaporin | Human-AQP2 | P41181 | N/A |
| Aquaporin | Mouse-AQP2 | P56402 | N/A |
| Aquaporin | Devil-AQP2 | ENSSHAT00000033099.1 | N/A |
| Aquaporin | Koala-AQP3 | ENSPCIT00000048802.2 | N/A |
| Aquaporin | Human-AQP3 | Q92482 | N/A |
| Aquaporin | Mouse-AQP3 | Q8R2N1 | N/A |
| Aquaporin | Opossum-AQP3 | ENSMODT00000004790.4 | N/A |
| Aquaporin | Devil-AQP3 | ENSSHAT00000003653.2 | N/A |
| Aquaporin | Koala-AQP4 | A0A6P5J0L6 | N/A |
| Aquaporin | Human-AQP4 | P55087 | N/A |
| Aquaporin | Mouse-AQP4 | P55088 | N/A |
| Aquaporin | Devil-AQP4 | ENSSHAT00000017782.2 | N/A |
| Aquaporin | Opossum-AQP4 | ENSMODT00000027360.4 | N/A |
| Aquaporin | Koala-AQP5 | A0A6P5IRY7 | N/A |
| Aquaporin | Human-AQP5 | P55064 | N/A |
| Aquaporin | Mouse-AQP5 | Q9WTY4 | N/A |
| Aquaporin | Opossum-AQP5 | ENSMODT00000000296.3 | N/A |
| Aquaporin | Devil-AQP5 | ENSSHAT00000004355.2 | N/A |
| Aquaporin | Human-AQP6 | Q13520 | N/A |
| Aquaporin | Mouse-AQP6 | Q8C4A0 | N/A |
| Aquaporin | Devil-AQP6 | ENSSHAT00000039717.1 | N/A |
| Aquaporin | Koala-AQP6 | A0A6P5IHT5 | N/A |
| Aquaporin | Koala-AQP7 | A0A6P5K9N9 | N/A |
| Aquaporin | Koala-AQP7 | A0A6P5JQG0 | N/A |
| Aquaporin | Human-AQP7 | O14520 | N/A |
| Aquaporin | Mouse-AQP7 | O54794 | N/A |
| Aquaporin | Koala-AQP8 | A0A6P5KZI2 | N/A |
| Aquaporin | Human-AQP8 | O94778 | N/A |
| Aquaporin | Mouse-AQP8 | P56404 | N/A |
| Aquaporin | Devil-AQP8 | ENSSHAT00000004706.2 | N/A |
| Aquaporin | Opossum-AQP8 | ENSMODT00000020737.3 | N/A |
| Aquaporin | Koala-AQP9 | A0A6P5JYG4 | N/A |
| Aquaporin | Human-AQP9 | O43315 | N/A |
| Aquaporin | Mouse-AQP9 | Q9JJJ3 | N/A |
| Aquaporin | Opossum-AQP9 | ENSMODT00000042732.2 | N/A |
| Aquaporin | Devil-AQP9 | ENSSHAT00000013494.2 | N/A |
| Aquaporin | Koala-AQP10 | ENSPCIT00000049244.2 | N/A |
| Aquaporin | Human-AQP10 | Q96PS8 | N/A |
| Aquaporin | Devil-AQP10 | ENSSHAT00000025526.1 | N/A |
| Aquaporin | Opossum-AQP10 | ENSMODT00000021823.3 | N/A |
| Aquaporin | Koala-AQP11 | ENSPCIT00000020412.2 | N/A |
| Aquaporin | Human-AQP11 | Q8NBQ7 | N/A |

| | | | |
|---|---|---|---|
| Aquaporin | Mouse-AQP11 | Q8BHH1 | N/A |
| Aquaporin | Opossum-AQP11 | ENSMODT00000006322.3 | N/A |
| Aquaporin | Devil-AQP11 | ENSSHAT00000044826.1 | N/A |
| Aquaporin | Koala-AQP12 | A0A6P5K9Q3 | N/A |
| Aquaporin | Human-AQP12B | A6NM10 | N/A |
| Aquaporin | Human-AQP12A | Q8IXF9 | N/A |
| Aquaporin | Mouse-AQP12 | Q8CHJ2 | N/A |
| Aquaporin | Opossum-AQP12-like | | N/A |
| V1R | monDomV1R1201_md | GCF_000002295.2 | Figure 6 |
| V1R | monDomV1R1202_md | GCF_000002295.2 | Figure 6 |
| V1R | monDomV1R1203_md | GCF_000002295.2 | Figure 6 |
| V1R | monDomV1R1204_md | GCF_000002295.2 | Figure 6 |
| V1R | monDomV1R1205_md | GCF_000002295.2 | Figure 6 |
| V1R | monDomV1R1206_md | GCF_000002295.2 | Figure 6 |
| V1R | monDomV1R1207_md | GCF_000002295.2 | Figure 6 |
| V1R | monDomV1R1208_md | GCF_000002295.2 | Figure 6 |
| V1R | monDomV1R1209_md | GCF_000002295.2 | Figure 6 |
| V1R | monDomV1R1210_md | GCF_000002295.2 | Figure 6 |
| V1R | monDomV1R1211_md | GCF_000002295.2 | Figure 6 |
| V1R | monDomV1R1212_md | GCF_000002295.2 | Figure 6 |
| V1R | monDomV1R1213_md | GCF_000002295.2 | Figure 6 |
| V1R | monDomV1R1214_md | GCF_000002295.2 | Figure 6 |
| V1R | monDomV1R1215_md | GCF_000002295.2 | Figure 6 |
| V1R | monDomV1R1216_md | GCF_000002295.2 | Figure 6 |
| V1R | monDomV1R1217_md | GCF_000002295.2 | Figure 6 |
| V1R | monDomV1R1218_md | GCF_000002295.2 | Figure 6 |
| V1R | monDomV1R1219_md | GCF_000002295.2 | Figure 6 |
| V1R | monDomV1R1220_md | GCF_000002295.2 | Figure 6 |
| V1R | monDomV1R1221_md | GCF_000002295.2 | Figure 6 |
| V1R | monDomV1R1222_md | GCF_000002295.2 | Figure 6 |
| V1R | monDomV1R1223_md | GCF_000002295.2 | Figure 6 |
| V1R | monDomV1R1224_md | GCF_000002295.2 | Figure 6 |
| V1R | monDomV1R1225_md | GCF_000002295.2 | Figure 6 |
| V1R | monDomV1R1226_md | GCF_000002295.2 | Figure 6 |
| V1R | monDomV1R1227_md | GCF_000002295.2 | Figure 6 |
| V1R | monDomV1R1228_md | GCF_000002295.2 | Figure 6 |
| V1R | monDomV1R1229_md | GCF_000002295.2 | Figure 6 |
| V1R | monDomV1R1230_md | GCF_000002295.2 | Figure 6 |
| V1R | monDomV1R1231_md | GCF_000002295.2 | Figure 6 |
| V1R | monDomV1R1232_md | GCF_000002295.2 | Figure 6 |
| V1R | monDomV1R1233_md | GCF_000002295.2 | Figure 6 |
| V1R | monDomV1R1234_md | GCF_000002295.2 | Figure 6 |
| V1R | monDomV1R1235_md | GCF_000002295.2 | Figure 6 |
| V1R | monDomV1R1236_md | GCF_000002295.2 | Figure 6 |
| V1R | monDomV1R1237_md | GCF_000002295.2 | Figure 6 |
| V1R | monDomV1R1238_md | GCF_000002295.2 | Figure 6 |
| V1R | monDomV1R1239_md | GCF_000002295.2 | Figure 6 |
| V1R | monDomV1R1240_md | GCF_000002295.2 | Figure 6 |
| V1R | monDomV1R1241_md | GCF_000002295.2 | Figure 6 |
| V1R | monDomV1R1242_md | GCF_000002295.2 | Figure 6 |

| | | | |
|---|---|---|---|
| V1R | monDomV1R1243_md | GCF_000002295.2 | Figure 6 |
| V1R | monDomV1R1244_md | GCF_000002295.2 | Figure 6 |
| V1R | monDomV1R1245_md | GCF_000002295.2 | Figure 6 |
| V1R | monDomV1R1246_md | GCF_000002295.2 | Figure 6 |
| V1R | monDomV1R1247_md | GCF_000002295.2 | Figure 6 |
| V1R | monDomV1R1248_md | GCF_000002295.2 | Figure 6 |
| V1R | monDomV1R1249_md | GCF_000002295.2 | Figure 6 |
| V1R | monDomV1R1250_md | GCF_000002295.2 | Figure 6 |
| V1R | monDomV1R1251_md | GCF_000002295.2 | Figure 6 |
| V1R | monDomV1R1252_md | GCF_000002295.2 | Figure 6 |
| V1R | monDomV1R1253_md | GCF_000002295.2 | Figure 6 |
| V1R | monDomV1R1254_md | GCF_000002295.2 | Figure 6 |
| V1R | monDomV1R1255_md | GCF_000002295.2 | Figure 6 |
| V1R | monDomV1R1256_md | GCF_000002295.2 | Figure 6 |
| V1R | monDomV1R1257_md | GCF_000002295.2 | Figure 6 |
| V1R | monDomV1R1258_md | GCF_000002295.2 | Figure 6 |
| V1R | monDomV1R1259_md | GCF_000002295.2 | Figure 6 |
| V1R | monDomV1R1260_md | GCF_000002295.2 | Figure 6 |
| V1R | monDomV1R1261_md | GCF_000002295.2 | Figure 6 |
| V1R | monDomV1R1262_md | GCF_000002295.2 | Figure 6 |
| V1R | monDomV1R1263_md | GCF_000002295.2 | Figure 6 |
| V1R | monDomV1R1264_md | GCF_000002295.2 | Figure 6 |
| V1R | monDomV1R1265_md | GCF_000002295.2 | Figure 6 |
| V1R | monDomV1R1266_md | GCF_000002295.2 | Figure 6 |
| V1R | monDomV1R1267_md | GCF_000002295.2 | Figure 6 |
| V1R | monDomV1R1268_md | GCF_000002295.2 | Figure 6 |
| V1R | monDomV1R1269_md | GCF_000002295.2 | Figure 6 |
| V1R | monDomV1R1270_md | GCF_000002295.2 | Figure 6 |
| V1R | monDomV1R1271_md | GCF_000002295.2 | Figure 6 |
| V1R | monDomV1R1272_md | GCF_000002295.2 | Figure 6 |
| V1R | monDomV1R1273_md | GCF_000002295.2 | Figure 6 |
| V1R | monDomV1R1274_md | GCF_000002295.2 | Figure 6 |
| V1R | monDomV1R1275_md | GCF_000002295.2 | Figure 6 |
| V1R | monDomV1R1276_md | GCF_000002295.2 | Figure 6 |
| V1R | monDomV1R1277_md | GCF_000002295.2 | Figure 6 |
| V1R | monDomV1R1278_md | GCF_000002295.2 | Figure 6 |
| V1R | monDomV1R1279_md | GCF_000002295.2 | Figure 6 |
| V1R | monDomV1R1280_md | GCF_000002295.2 | Figure 6 |
| V1R | monDomV1R1281_md | GCF_000002295.2 | Figure 6 |
| V1R | monDomV1R1282_md | GCF_000002295.2 | Figure 6 |
| V1R | monDomV1R1283_md | GCF_000002295.2 | Figure 6 |
| V1R | monDomV1R1284_md | GCF_000002295.2 | Figure 6 |
| V1R | monDomV1R1285_md | GCF_000002295.2 | Figure 6 |
| V1R | monDomV1R1286_md | GCF_000002295.2 | Figure 6 |
| V1R | monDomV1R1287_md | GCF_000002295.2 | Figure 6 |
| V1R | monDomV1R1288_md | GCF_000002295.2 | Figure 6 |
| V1R | monDomV1R1289_md | GCF_000002295.2 | Figure 6 |
| V1R | monDomV1R1290_md | GCF_000002295.2 | Figure 6 |
| V1R | monDomV1R1291_md | GCF_000002295.2 | Figure 6 |
| V1R | monDomV1R1292_md | GCF_000002295.2 | Figure 6 |

| | | | |
|---|---|---|---|
| V1R | monDomV1R1293_md | GCF_000002295.2 | Figure 6 |
| V1R | monDomV1R1294_md | GCF_000002295.2 | Figure 6 |
| V1R | monDomV1R1295_md | GCF_000002295.2 | Figure 6 |
| V1R | MumuV1R107_mm | D3YTY1 | Figure 6 |
| V1R | MumuV1R13_mm | G5E8I3 | Figure 6 |
| V1R | MumuV1R189_mm | Q8K3N3 | Figure 6 |
| V1R | MumuV1R201_mm | Q8R262 | Figure 6 |
| V1R | MumuV1R210_mm | Q8R274 | Figure 6 |
| V1R | MumuV1R214_mm | Q8R279 | Figure 6 |
| V1R | MumuV1R217_mm | Q8R270 | Figure 6 |
| V1R | MumuV1R218_mm | Q8R261 | Figure 6 |
| V1R | MumuV1R29_mm | Q9EQ41 | Figure 6 |
| V1R | MumuV1R30_mm | Q8R2D2 | Figure 6 |
| V1R | MumuV1R37_mm | A0A2I3BPJ6 | Figure 6 |
| V1R | MumuV1R3_mm | A2AMT7 | Figure 6 |
| V1R | MumuV1R40_mm | Q9EQ46 | Figure 6 |
| V1R | MumuV1R41_mm | Q9EQ44 | Figure 6 |
| V1R | MumuV1R42_mm | Q8VBS7 | Figure 6 |
| V1R | MumuV1R43_mm | Q8VIC9 | Figure 6 |
| V1R | MumuV1R44_mm | Q9EQ44 | Figure 6 |
| V1R | MumuV1R45_mm | Q8VIC7 | Figure 6 |
| V1R | MumuV1R46_mm | Q9EQ45 | Figure 6 |
| V1R | MumuV1R47_mm | Q9EQ51 | Figure 6 |
| V1R | MumuV1R48_mm | Q9EQ52 | Figure 6 |
| V1R | MumuV1R49_mm | Q9WUF1 | Figure 6 |
| V1R | MumuV1R50_mm | Q9EP51 | Figure 6 |
| V1R | MumuV1R51_mm | Q8VIC6 | Figure 6 |
| V1R | MumuV1R52_mm | Q9EP79 | Figure 6 |
| V1R | MumuV1R53_mm | Q9EP93 | Figure 6 |
| V1R | MumuV1R54_mm | Q9EPB8 | Figure 6 |
| V1R | MumuV1R69_mm | Q8VIC1 | Figure 6 |
| V1R | MumuV1R71_mm | Q8VIC0 | Figure 6 |
| V1R | MumuV1R75_mm | Q8R289 | Figure 6 |
| V1R | MumuV1R87_mm | Q8R255 | Figure 6 |
| V1R | MumuV1RA11_mm | Q8R2E6 | Figure 6 |
| V1R | MumuV1RA8_mm | Q9EQ48 | Figure 6 |
| V1R | OranV1R106.1_oa | GCA_004115215.4 | Figure 6 |
| V1R | OranV1R11.1_oa | GCA_004115215.4 | Figure 6 |
| V1R | OranV1R111.1_oa | GCA_004115215.4 | Figure 6 |
| V1R | OranV1R114.1_oa | GCA_004115215.4 | Figure 6 |
| V1R | OranV1R115.1_oa | GCA_004115215.4 | Figure 6 |
| V1R | OranV1R120.1_oa | GCA_004115215.4 | Figure 6 |
| V1R | OranV1R126.1_oa | GCA_004115215.4 | Figure 6 |
| V1R | OranV1R128.1_oa | GCA_004115215.4 | Figure 6 |
| V1R | OranV1R133.1_oa | GCA_004115215.4 | Figure 6 |
| V1R | OranV1R139.1_oa | GCA_004115215.4 | Figure 6 |
| V1R | OranV1R141.1_oa | GCA_004115215.4 | Figure 6 |
| V1R | OranV1R146.1_oa | GCA_004115215.4 | Figure 6 |
| V1R | OranV1R148.1_oa | GCA_004115215.4 | Figure 6 |
| V1R | OranV1R15.1_oa | GCA_004115215.4 | Figure 6 |

| | | | |
|---|---|---|---|
| V1R | OranV1R152.1_oa | GCA_004115215.4 | Figure 6 |
| V1R | OranV1R153.1_oa | GCA_004115215.4 | Figure 6 |
| V1R | OranV1R158.1_oa | GCA_004115215.4 | Figure 6 |
| V1R | OranV1R160.1_oa | GCA_004115215.4 | Figure 6 |
| V1R | OranV1R163.1_oa | GCA_004115215.4 | Figure 6 |
| V1R | OranV1R166.1_oa | GCA_004115215.4 | Figure 6 |
| V1R | OranV1R169.1_oa | GCA_004115215.4 | Figure 6 |
| V1R | OranV1R170.1_oa | GCA_004115215.4 | Figure 6 |
| V1R | OranV1R175.1_oa | GCA_004115215.4 | Figure 6 |
| V1R | OranV1R177.1_oa | GCA_004115215.4 | Figure 6 |
| V1R | OranV1R179.1_oa | GCA_004115215.4 | Figure 6 |
| V1R | OranV1R180.1_oa | GCA_004115215.4 | Figure 6 |
| V1R | OranV1R181.1_oa | GCA_004115215.4 | Figure 6 |
| V1R | OranV1R185.1_oa | GCA_004115215.4 | Figure 6 |
| V1R | OranV1R188.1_oa | GCA_004115215.4 | Figure 6 |
| V1R | OranV1R205.1_oa | GCA_004115215.4 | Figure 6 |
| V1R | OranV1R214.1_oa | GCA_004115215.4 | Figure 6 |
| V1R | OranV1R216.1_oa | GCA_004115215.4 | Figure 6 |
| V1R | OranV1R218.1_oa | GCA_004115215.4 | Figure 6 |
| V1R | OranV1R219.1_oa | GCA_004115215.4 | Figure 6 |
| V1R | OranV1R223.1_oa | GCA_004115215.4 | Figure 6 |
| V1R | OranV1R224.1_oa | GCA_004115215.4 | Figure 6 |
| V1R | OranV1R227.1_oa | GCA_004115215.4 | Figure 6 |
| V1R | OranV1R231.1_oa | GCA_004115215.4 | Figure 6 |
| V1R | OranV1R246.1_oa | GCA_004115215.4 | Figure 6 |
| V1R | OranV1R250.1_oa | GCA_004115215.4 | Figure 6 |
| V1R | OranV1R263.1_oa | GCA_004115215.4 | Figure 6 |
| V1R | OranV1R265.1_oa | GCA_004115215.4 | Figure 6 |
| V1R | OranV1R267.1_oa | GCA_004115215.4 | Figure 6 |
| V1R | OranV1R268.1_oa | GCA_004115215.4 | Figure 6 |
| V1R | OranV1R270.1_oa | GCA_004115215.4 | Figure 6 |
| V1R | OranV1R272.1_oa | GCA_004115215.4 | Figure 6 |
| V1R | OranV1R278.1_oa | GCA_004115215.4 | Figure 6 |
| V1R | OranV1R280.1_oa | GCA_004115215.4 | Figure 6 |
| V1R | OranV1R281.1_oa | GCA_004115215.4 | Figure 6 |
| V1R | OranV1R282.1_oa | GCA_004115215.4 | Figure 6 |
| V1R | OranV1R284.1_oa | GCA_004115215.4 | Figure 6 |
| V1R | OranV1R285.1_oa | GCA_004115215.4 | Figure 6 |
| V1R | OranV1R286.1_oa | GCA_004115215.4 | Figure 6 |
| V1R | OranV1R287.1_oa | GCA_004115215.4 | Figure 6 |
| V1R | OranV1R288.1_oa | GCA_004115215.4 | Figure 6 |
| V1R | OranV1R289.1_oa | GCA_004115215.4 | Figure 6 |
| V1R | OranV1R290.1_oa | GCA_004115215.4 | Figure 6 |
| V1R | OranV1R291.1_oa | GCA_004115215.4 | Figure 6 |
| V1R | OranV1R292.1_oa | GCA_004115215.4 | Figure 6 |
| V1R | OranV1R293.1_oa | GCA_004115215.4 | Figure 6 |
| V1R | OranV1R294.1_oa | GCA_004115215.4 | Figure 6 |
| V1R | OranV1R295.1_oa | GCA_004115215.4 | Figure 6 |
| V1R | OranV1R296.1_oa | GCA_004115215.4 | Figure 6 |
| V1R | OranV1R297.1_oa | GCA_004115215.4 | Figure 6 |

| | | | |
|---|---|---|---|
| V1R | OranV1R298.1_oa | GCA_004115215.4 | Figure 6 |
| V1R | OranV1R299.1_oa | GCA_004115215.4 | Figure 6 |
| V1R | OranV1R30.1_oa | GCA_004115215.4 | Figure 6 |
| V1R | OranV1R300.1_oa | GCA_004115215.4 | Figure 6 |
| V1R | OranV1R301.1_oa | GCA_004115215.4 | Figure 6 |
| V1R | OranV1R306.1_oa | GCA_004115215.4 | Figure 6 |
| V1R | OranV1R307.1_oa | GCA_004115215.4 | Figure 6 |
| V1R | OranV1R309.1_oa | GCA_004115215.4 | Figure 6 |
| V1R | OranV1R31.1_oa | GCA_004115215.4 | Figure 6 |
| V1R | OranV1R310.1_oa | GCA_004115215.4 | Figure 6 |
| V1R | OranV1R311.1_oa | GCA_004115215.4 | Figure 6 |
| V1R | OranV1R312.1_oa | GCA_004115215.4 | Figure 6 |
| V1R | OranV1R313.1_oa | GCA_004115215.4 | Figure 6 |
| V1R | OranV1R314.1_oa | GCA_004115215.4 | Figure 6 |
| V1R | OranV1R316.1_oa | GCA_004115215.4 | Figure 6 |
| V1R | OranV1R318.1_oa | GCA_004115215.4 | Figure 6 |
| V1R | OranV1R321.1_oa | GCA_004115215.4 | Figure 6 |
| V1R | OranV1R322.1_oa | GCA_004115215.4 | Figure 6 |
| V1R | OranV1R323.1_oa | GCA_004115215.4 | Figure 6 |
| V1R | OranV1R324.1_oa | GCA_004115215.4 | Figure 6 |
| V1R | OranV1R325.1_oa | GCA_004115215.4 | Figure 6 |
| V1R | OranV1R326.1_oa | GCA_004115215.4 | Figure 6 |
| V1R | OranV1R327.1_oa | GCA_004115215.4 | Figure 6 |
| V1R | OranV1R328.1_oa | GCA_004115215.4 | Figure 6 |
| V1R | OranV1R329.1_oa | GCA_004115215.4 | Figure 6 |
| V1R | OranV1R330.1_oa | GCA_004115215.4 | Figure 6 |
| V1R | OranV1R334.1_oa | GCA_004115215.4 | Figure 6 |
| V1R | OranV1R336.1_oa | GCA_004115215.4 | Figure 6 |
| V1R | OranV1R337.1_oa | GCA_004115215.4 | Figure 6 |
| V1R | OranV1R339.1_oa | GCA_004115215.4 | Figure 6 |
| V1R | OranV1R34.1_oa | GCA_004115215.4 | Figure 6 |
| V1R | OranV1R340.1_oa | GCA_004115215.4 | Figure 6 |
| V1R | OranV1R342.1_oa | GCA_004115215.4 | Figure 6 |
| V1R | OranV1R343.1_oa | GCA_004115215.4 | Figure 6 |
| V1R | OranV1R344.1_oa | GCA_004115215.4 | Figure 6 |
| V1R | OranV1R345.1_oa | GCA_004115215.4 | Figure 6 |
| V1R | OranV1R35.1_oa | GCA_004115215.4 | Figure 6 |
| V1R | OranV1R350.1_oa | GCA_004115215.4 | Figure 6 |
| V1R | OranV1R351.1_oa | GCA_004115215.4 | Figure 6 |
| V1R | OranV1R353.1_oa | GCA_004115215.4 | Figure 6 |
| V1R | OranV1R355.1_oa | GCA_004115215.4 | Figure 6 |
| V1R | OranV1R359.1_oa | GCA_004115215.4 | Figure 6 |
| V1R | OranV1R36.1_oa | GCA_004115215.4 | Figure 6 |
| V1R | OranV1R360.1_oa | GCA_004115215.4 | Figure 6 |
| V1R | OranV1R362.1_oa | GCA_004115215.4 | Figure 6 |
| V1R | OranV1R368.1_oa | GCA_004115215.4 | Figure 6 |
| V1R | OranV1R37.1_oa | GCA_004115215.4 | Figure 6 |
| V1R | OranV1R370.1_oa | GCA_004115215.4 | Figure 6 |
| V1R | OranV1R386.1_oa | GCA_004115215.4 | Figure 6 |
| V1R | OranV1R392.1_oa | GCA_004115215.4 | Figure 6 |

| | | | |
|---|---|---|---|
| V1R | OranV1R409.1_oa | GCA_004115215.4 | Figure 6 |
| V1R | OranV1R41.1_oa | GCA_004115215.4 | Figure 6 |
| V1R | OranV1R42.1_oa | GCA_004115215.4 | Figure 6 |
| V1R | OranV1R421.1_oa | GCA_004115215.4 | Figure 6 |
| V1R | OranV1R43.1_oa | GCA_004115215.4 | Figure 6 |
| V1R | OranV1R430.1_oa | GCA_004115215.4 | Figure 6 |
| V1R | OranV1R432.1_oa | GCA_004115215.4 | Figure 6 |
| V1R | OranV1R433.1_oa | GCA_004115215.4 | Figure 6 |
| V1R | OranV1R434.1_oa | GCA_004115215.4 | Figure 6 |
| V1R | OranV1R435.1_oa | GCA_004115215.4 | Figure 6 |
| V1R | OranV1R436.1_oa | GCA_004115215.4 | Figure 6 |
| V1R | OranV1R437.1_oa | GCA_004115215.4 | Figure 6 |
| V1R | OranV1R438.1_oa | GCA_004115215.4 | Figure 6 |
| V1R | OranV1R439.1_oa | GCA_004115215.4 | Figure 6 |
| V1R | OranV1R440.1_oa | GCA_004115215.4 | Figure 6 |
| V1R | OranV1R441.1_oa | GCA_004115215.4 | Figure 6 |
| V1R | OranV1R442.1_oa | GCA_004115215.4 | Figure 6 |
| V1R | OranV1R445.1_oa | GCA_004115215.4 | Figure 6 |
| V1R | OranV1R446.1_oa | GCA_004115215.4 | Figure 6 |
| V1R | OranV1R447.1_oa | GCA_004115215.4 | Figure 6 |
| V1R | OranV1R448.1_oa | GCA_004115215.4 | Figure 6 |
| V1R | OranV1R449.1_oa | GCA_004115215.4 | Figure 6 |
| V1R | OranV1R45.1_oa | GCA_004115215.4 | Figure 6 |
| V1R | OranV1R450.1_oa | GCA_004115215.4 | Figure 6 |
| V1R | OranV1R451.1_oa | GCA_004115215.4 | Figure 6 |
| V1R | OranV1R453.1_oa | GCA_004115215.4 | Figure 6 |
| V1R | OranV1R454.1_oa | GCA_004115215.4 | Figure 6 |
| V1R | OranV1R455.1_oa | GCA_004115215.4 | Figure 6 |
| V1R | OranV1R456.1_oa | GCA_004115215.4 | Figure 6 |
| V1R | OranV1R457.1_oa | GCA_004115215.4 | Figure 6 |
| V1R | OranV1R46.1_oa | GCA_004115215.4 | Figure 6 |
| V1R | OranV1R461.1_oa | GCA_004115215.4 | Figure 6 |
| V1R | OranV1R462.1_oa | GCA_004115215.4 | Figure 6 |
| V1R | OranV1R463.1_oa | GCA_004115215.4 | Figure 6 |
| V1R | OranV1R464.1_oa | GCA_004115215.4 | Figure 6 |
| V1R | OranV1R465.1_oa | GCA_004115215.4 | Figure 6 |
| V1R | OranV1R466.1_oa | GCA_004115215.4 | Figure 6 |
| V1R | OranV1R467.1_oa | GCA_004115215.4 | Figure 6 |
| V1R | OranV1R468.1_oa | GCA_004115215.4 | Figure 6 |
| V1R | OranV1R469.1_oa | GCA_004115215.4 | Figure 6 |
| V1R | OranV1R47.1_oa | GCA_004115215.4 | Figure 6 |
| V1R | OranV1R470.1_oa | GCA_004115215.4 | Figure 6 |
| V1R | OranV1R473.1_oa | GCA_004115215.4 | Figure 6 |
| V1R | OranV1R474.1_oa | GCA_004115215.4 | Figure 6 |
| V1R | OranV1R475.1_oa | GCA_004115215.4 | Figure 6 |
| V1R | OranV1R476.1_oa | GCA_004115215.4 | Figure 6 |
| V1R | OranV1R477.1_oa | GCA_004115215.4 | Figure 6 |
| V1R | OranV1R478.1_oa | GCA_004115215.4 | Figure 6 |
| V1R | OranV1R479.1_oa | GCA_004115215.4 | Figure 6 |
| V1R | OranV1R48.1_oa | GCA_004115215.4 | Figure 6 |

| | | | |
|---|---|---|---|
| V1R | OranV1R480.1_oa | GCA_004115215.4 | Figure 6 |
| V1R | OranV1R481.1_oa | GCA_004115215.4 | Figure 6 |
| V1R | OranV1R482.1_oa | GCA_004115215.4 | Figure 6 |
| V1R | OranV1R483.1_oa | GCA_004115215.4 | Figure 6 |
| V1R | OranV1R485.1_oa | GCA_004115215.4 | Figure 6 |
| V1R | OranV1R488.1_oa | GCA_004115215.4 | Figure 6 |
| V1R | OranV1R50.1_oa | GCA_004115215.4 | Figure 6 |
| V1R | OranV1R500.1_oa | GCA_004115215.4 | Figure 6 |
| V1R | OranV1R51.1_oa | GCA_004115215.4 | Figure 6 |
| V1R | OranV1R510.1_oa | GCA_004115215.4 | Figure 6 |
| V1R | OranV1R516.1_oa | GCA_004115215.4 | Figure 6 |
| V1R | OranV1R517.1_oa | GCA_004115215.4 | Figure 6 |
| V1R | OranV1R52.1_oa | GCA_004115215.4 | Figure 6 |
| V1R | OranV1R548.1_oa | GCA_004115215.4 | Figure 6 |
| V1R | OranV1R563.1_oa | GCA_004115215.4 | Figure 6 |
| V1R | OranV1R564.1_oa | GCA_004115215.4 | Figure 6 |
| V1R | OranV1R565.1_oa | GCA_004115215.4 | Figure 6 |
| V1R | OranV1R567.1_oa | GCA_004115215.4 | Figure 6 |
| V1R | OranV1R571.1_oa | GCA_004115215.4 | Figure 6 |
| V1R | OranV1R576.1_oa | GCA_004115215.4 | Figure 6 |
| V1R | OranV1R58.1_oa | GCA_004115215.4 | Figure 6 |
| V1R | OranV1R586.1_oa | GCA_004115215.4 | Figure 6 |
| V1R | OranV1R587.1_oa | GCA_004115215.4 | Figure 6 |
| V1R | OranV1R598.1_oa | GCA_004115215.4 | Figure 6 |
| V1R | OranV1R60.1_oa | GCA_004115215.4 | Figure 6 |
| V1R | OranV1R600.1_oa | GCA_004115215.4 | Figure 6 |
| V1R | OranV1R610.1_oa | GCA_004115215.4 | Figure 6 |
| V1R | OranV1R619.1_oa | GCA_004115215.4 | Figure 6 |
| V1R | OranV1R62.1_oa | GCA_004115215.4 | Figure 6 |
| V1R | OranV1R626.1_oa | GCA_004115215.4 | Figure 6 |
| V1R | OranV1R63.1_oa | GCA_004115215.4 | Figure 6 |
| V1R | OranV1R634.1_oa | GCA_004115215.4 | Figure 6 |
| V1R | OranV1R639.1_oa | GCA_004115215.4 | Figure 6 |
| V1R | OranV1R644.1_oa | GCA_004115215.4 | Figure 6 |
| V1R | OranV1R649.1_oa | GCA_004115215.4 | Figure 6 |
| V1R | OranV1R65.1_oa | GCA_004115215.4 | Figure 6 |
| V1R | OranV1R651.1_oa | GCA_004115215.4 | Figure 6 |
| V1R | OranV1R656.1_oa | GCA_004115215.4 | Figure 6 |
| V1R | OranV1R660.1_oa | GCA_004115215.4 | Figure 6 |
| V1R | OranV1R661.1_oa | GCA_004115215.4 | Figure 6 |
| V1R | OranV1R665.1_oa | GCA_004115215.4 | Figure 6 |
| V1R | OranV1R67.1_oa | GCA_004115215.4 | Figure 6 |
| V1R | OranV1R674.1_oa | GCA_004115215.4 | Figure 6 |
| V1R | OranV1R68.1_oa | GCA_004115215.4 | Figure 6 |
| V1R | OranV1R685.1_oa | GCA_004115215.4 | Figure 6 |
| V1R | OranV1R693.1_oa | GCA_004115215.4 | Figure 6 |
| V1R | OranV1R699.1_oa | GCA_004115215.4 | Figure 6 |
| V1R | OranV1R71.1_oa | GCA_004115215.4 | Figure 6 |
| V1R | OranV1R72.1_oa | GCA_004115215.4 | Figure 6 |
| V1R | OranV1R723.1_oa | GCA_004115215.4 | Figure 6 |

| | | | |
|---|---|---|---|
| V1R | OranV1R725.1_oa | GCA_004115215.4 | Figure 6 |
| V1R | OranV1R73.1_oa | GCA_004115215.4 | Figure 6 |
| V1R | OranV1R731.1_oa | GCA_004115215.4 | Figure 6 |
| V1R | OranV1R732.1_oa | GCA_004115215.4 | Figure 6 |
| V1R | OranV1R734.1_oa | GCA_004115215.4 | Figure 6 |
| V1R | OranV1R736.1_oa | GCA_004115215.4 | Figure 6 |
| V1R | OranV1R748.1_oa | GCA_004115215.4 | Figure 6 |
| V1R | OranV1R75.1_oa | GCA_004115215.4 | Figure 6 |
| V1R | OranV1R753.1_oa | GCA_004115215.4 | Figure 6 |
| V1R | OranV1R76.1_oa | GCA_004115215.4 | Figure 6 |
| V1R | OranV1R767.1_oa | GCA_004115215.4 | Figure 6 |
| V1R | OranV1R773.1_oa | GCA_004115215.4 | Figure 6 |
| V1R | OranV1R775.1_oa | GCA_004115215.4 | Figure 6 |
| V1R | OranV1R780.1_oa | GCA_004115215.4 | Figure 6 |
| V1R | OranV1R782.1_oa | GCA_004115215.4 | Figure 6 |
| V1R | OranV1R784.1_oa | GCA_004115215.4 | Figure 6 |
| V1R | OranV1R788.1_oa | GCA_004115215.4 | Figure 6 |
| V1R | OranV1R790.1_oa | GCA_004115215.4 | Figure 6 |
| V1R | OranV1R793.1_oa | GCA_004115215.4 | Figure 6 |
| V1R | OranV1R799.1_oa | GCA_004115215.4 | Figure 6 |
| V1R | OranV1R80.1_oa | GCA_004115215.4 | Figure 6 |
| V1R | OranV1R802.1_oa | GCA_004115215.4 | Figure 6 |
| V1R | OranV1R810.1_oa | GCA_004115215.4 | Figure 6 |
| V1R | OranV1R813.1_oa | GCA_004115215.4 | Figure 6 |
| V1R | OranV1R820.1_oa | GCA_004115215.4 | Figure 6 |
| V1R | OranV1R822.1_oa | GCA_004115215.4 | Figure 6 |
| V1R | OranV1R827.1_oa | GCA_004115215.4 | Figure 6 |
| V1R | OranV1R832.1_oa | GCA_004115215.4 | Figure 6 |
| V1R | OranV1R841.1_oa | GCA_004115215.4 | Figure 6 |
| V1R | OranV1R842.1_oa | GCA_004115215.4 | Figure 6 |
| V1R | OranV1R857.1_oa | GCA_004115215.4 | Figure 6 |
| V1R | OranV1R860.1_oa | GCA_004115215.4 | Figure 6 |
| V1R | OranV1R861.1_oa | GCA_004115215.4 | Figure 6 |
| V1R | OranV1R866.1_oa | GCA_004115215.4 | Figure 6 |
| V1R | OranV1R867.1_oa | GCA_004115215.4 | Figure 6 |
| V1R | OranV1R868.1_oa | GCA_004115215.4 | Figure 6 |
| V1R | OranV1R874.1_oa | GCA_004115215.4 | Figure 6 |
| V1R | OranV1R875.1_oa | GCA_004115215.4 | Figure 6 |
| V1R | OranV1R878.1_oa | GCA_004115215.4 | Figure 6 |
| V1R | OranV1R88.1_oa | GCA_004115215.4 | Figure 6 |
| V1R | OranV1R881.1_oa | GCA_004115215.4 | Figure 6 |
| V1R | OranV1R882.1_oa | GCA_004115215.4 | Figure 6 |
| V1R | OranV1R883.1_oa | GCA_004115215.4 | Figure 6 |
| V1R | OranV1R890.1_oa | GCA_004115215.4 | Figure 6 |
| V1R | OranV1R91.1_oa | GCA_004115215.4 | Figure 6 |
| V1R | OranV1R92.1_oa | GCA_004115215.4 | Figure 6 |
| V1R | OranV1R94.1_oa | GCA_004115215.4 | Figure 6 |
| V1R | OranV1R96.1_oa | GCA_004115215.4 | Figure 6 |
| V1R | TaacV1R1008.1_ta | GCA_015852505.1 | Figure 6 |
| V1R | TaacV1R1037.1_ta | GCA_015852505.1 | Figure 6 |

| | | | |
|---|---|---|---|
| V1R | TaacV1R119.1_ta | GCA_015852505.1 | Figure 6 |
| V1R | TaacV1R1219.1_ta | GCA_015852505.1 | Figure 6 |
| V1R | TaacV1R1239.1_ta | GCA_015852505.1 | Figure 6 |
| V1R | TaacV1R13.1_ta | GCA_015852505.1 | Figure 6 |
| V1R | TaacV1R142.1_ta | GCA_015852505.1 | Figure 6 |
| V1R | TaacV1R196.1_ta | GCA_015852505.1 | Figure 6 |
| V1R | TaacV1R363.1_ta | GCA_015852505.1 | Figure 6 |
| V1R | TaacV1R392.1_ta | GCA_015852505.1 | Figure 6 |
| V1R | TaacV1R397.1_ta | GCA_015852505.1 | Figure 6 |
| V1R | TaacV1R404.1_ta | GCA_015852505.1 | Figure 6 |
| V1R | TaacV1R469.1_ta | GCA_015852505.1 | Figure 6 |
| V1R | TaacV1R472.1_ta | GCA_015852505.1 | Figure 6 |
| V1R | TaacV1R503.1_ta | GCA_015852505.1 | Figure 6 |
| V1R | TaacV1R574.1_ta | GCA_015852505.1 | Figure 6 |
| V1R | TaacV1R578.1_ta | GCA_015852505.1 | Figure 6 |
| V1R | TaacV1R586.1_ta | GCA_015852505.1 | Figure 6 |
| V1R | TaacV1R59.1_ta | GCA_015852505.1 | Figure 6 |
| V1R | TaacV1R613.1_ta | GCA_015852505.1 | Figure 6 |
| V1R | TaacV1R615.1_ta | GCA_015852505.1 | Figure 6 |
| V1R | TaacV1R66.1_ta | GCA_015852505.1 | Figure 6 |
| V1R | TaacV1R864.1_ta | GCA_015852505.1 | Figure 6 |
| V1R | TaacV1R867.1_ta | GCA_015852505.1 | Figure 6 |
| V1R | TaacV1R9.1_ta | GCA_015852505.1 | Figure 6 |
| V1R | TaacV1R938.1_ta | GCA_015852505.1 | Figure 6 |
| V1R | TaacV1R986.1_ta | GCA_015852505.1 | Figure 6 |
| V1R | TaacV1R988.1_ta | GCA_015852505.1 | Figure 6 |
| V1R | Phci-VN1R1 | XP_020831561.1 | Figure 6 |
| V1R | Phci-VN1R2 | XP_020833384.1 | Figure 6 |
| V1R | Phci-VN1R3 | XP_020865409.1 | Figure 6 |
| V1R | Phci-VN1R4 | XP_020865367.1 | Figure 6 |
| V1R | HosaV1R1 | Q9GZP7 | Figure 6 |
| V1R | HosaV1R2 | Q8NFZ6 | Figure 6 |
| V1R | HosaV1R3 | Q9BXE9 | Figure 6 |
| V1R | HosaV1R4 | Q7Z5H5 | Figure 6 |
| V1R | HosaV1R5 | Q7Z5H4 | Figure 6 |
| V2R | ModoV2R653 | NP_001095921.1 | N/A |
| V2R | ModoV2R598 | NP_001093048.1 | N/A |
| V2R | ModoV2R615 | NP_001093039.1 | N/A |
| V2R | ModoV2R616 | NP_001093040.1 | N/A |
| V2R | ModoV2R657 | NP_001093066.1 | N/A |
| V2R | ModoV2R551 | NP_001093031.1 | N/A |
| V2R | MumuV2R116 | E9Q6I0 | N/A |
| V2R | MumuV2R116 | O70410 | N/A |
| V2R | MumuV2R26 | Q6TAC4 | N/A |
| V2R | PhciV2R26-like | XP_020834486.1 | N/A |
| V2R | Saha-V2R26-like | XP_031804987.1 | N/A |
| V2R | VourV2R26-like | XP_027702042.1 | N/A |
