## Supplementary Table 2 for "Genome assembly of the numbat (*Myrmecobius fasciatus*), the only termitivorous marsupial"

| **Supplementary Table 2.** Genomic coordinates of manually annotated taste and vomeronasal receptor genes and aquaporin genes in the numbat genome. p indicates potential pseudogene | | | | | |
|---|---|---|---|---|---|
| **Gene family** | **Gene name** | **Scaffold** | **Start** | **Stop** | **Strand** |
| TS1R | TAS1R1 | pri6555_MYRFA_USYD1HAP6555 | 148242 | 156185 | + |
| TS1R | TAS1R2 | pri1912_MYRFA_USYD1HAP1912 | 44829 | 55345 | + |
| TS1R | TAS1R3 | pri5031_MYRFA_USYD2HAP5031 | 63826 | 74159 | + |
| TS2R | TAS2R1 | pri304234_MYRFA_USYD1HAP304234 | 32054 | 32902 | - |
| TS2R | TAS2R723 | pri3565_MYRFA_USYD1HAP3565 | 168027 | 167155 | - |
| TS2R | TAS2R725 | pri3565_MYRFA_USYD1HAP3565 | 189028 | 188153 | - |
| TS2R | TAS2R729 | pri3565_MYRFA_USYD1HAP3565 | 245096 | 244152 | - |
| TS2R | TAS2R711 | pri4066_MYRFA_USYD2HAP4066 | 25733 | 26650 | + |
| TS2R | TAS2R4 | pri4066_MYRFA_USYD2HAP4066 | 35306 | 34467 | |
| TS2R | TAS2R714 | pri4066_MYRFA_USYD2HAP4066 | 96778 | 97662 | + |
| TS2R | TAS2R705A | pri6209_MYRFA_USYD1HAP6209 | 261346 | 260450 | - |
| TS2R | TAS2R41A | pri6209_MYRFA_USYD1HAP6209 | 281506 | 280556 | - |
| TS2R | TAS2R60 | pri6638_MYRFA_USYD1HAP6638 | 123638 | 124537 | + |
| TS2R | TAS2R702T | pri897_MYRFA_USYD2HAP897 | 499387 | 500010 | + |
| TS2R | TAS2R720p | pri3565_MYRFA_USYD1HAP3565 | 176123 | 175743 | - |
| TS2R | TAS2R726p | pri3565_MYRFA_USYD1HAP3565 | 218671 | 217753 | - |
| TS2R | TAS2R727p | pri3565_MYRFA_USYD1HAP3565 | 227927 | 227019 | - |
| TS2R | TAS2R728p | pri3565_MYRFA_USYD1HAP3565 | 261350 | 260625 | - |
| TS2R | TAS2R701p | pri4066_MYRFA_USYD2HAP4066 | 31675 | 31001 | - |
| TS2R | TAS2R710p | pri4066_MYRFA_USYD2HAP4066 | 73512 | 74420 | + |
| TS2R | TAS2R712p | pri4066_MYRFA_USYD2HAP4066 | 83993 | 84894 | + |
| TS2R | TAS2R713p | pri4066_MYRFA_USYD2HAP4066 | 92267 | 92720 | + |
| TS2R | TAS2R41Bp | pri6209_MYRFA_USYD1HAP6209 | 241349 | 240555 | - |
| TS2R | TAS2R705Bp | pri6209_MYRFA_USYD1HAP6209 | 297189 | 296773 | - |
| TS2R | TAS2R703p | pri897_MYRFA_USYD2HAP897 | 524783 | 525748 | + |
| VN1 | V1R1 | pri1719_MYRFA_USYD1HAP1719 | 120410 | 119523 | - |
| VN1 | V1R2 | pri204403_MYRFA_USYD1HAP204403 | 1456 | 539 | - |
| VN1 | V1R3 | pri220996_MYRFA_USYD1HAP220996 | 1548 | 2411 | + |
| VN1 | V1R4 | pri253177_MYRFA_USYD1HAP253177 | 1863 | 979 | - |
| VN1 | V1R5 | pri257492_MYRFA_USYD1HAP257492 | 51 | 908 | + |
| VN1 | V1R6 | pri262335_MYRFA_USYD1HAP262335 | 1647 | 2519 | + |
| VN1 | V1R7 | pri265445_MYRFA_USYD1HAP265445 | 10219 | 11196 | + |
| VN1 | V1R8 | pri274752_MYRFA_USYD1HAP274752 | 5190 | 6140 | + |
| VN1 | V1R9 | pri284998_MYRFA_USYD1HAP284998 | 8065 | 9009 | + |
| VN1 | V1R10 | pri285085_MYRFA_USYD1HAP285085 | 9392 | 8553 | - |
| VN1 | V1R11 | pri289009_MYRFA_USYD1HAP289009 | 7479 | 8405 | + |
| VN1 | V1R12 | pri290716_MYRFA_USYD1HAP290716 | 6510 | 7412 | + |
| VN1 | V1R13 | pri290767_MYRFA_USYD1HAP290767 | 810 | 1709 | + |
| VN1 | V1R14 | pri290767_MYRFA_USYD1HAP290767 | 27933 | 27001 | - |
| VN1 | V1R15 | pri290834_MYRFA_USYD1HAP290834 | 539 | 1423 | + |
| VN1 | V1R16 | pri295589_MYRFA_USYD1HAP295589 | 20782 | 19976 | - |
| VN1 | V1R17 | pri295817_MYRFA_USYD1HAP295817 | 4220 | 5140 | + |
| VN1 | V1R18 | pri296644_MYRFA_USYD1HAP296644 | 7957 | 7007 | - |
| VN1 | V1R19 | pri297495_MYRFA_USYD1HAP297495 | 7680 | 8567 | + |
| VN1 | V1R20 | pri298039_MYRFA_USYD1HAP298039 | 4153 | 3344 | - |
| VN1 | V1R21 | pri298441_MYRFA_USYD1HAP298441 | 14754 | 13894 | - |
| VN1 | V1R22 | pri299312_MYRFA_USYD1HAP299312 | 1499 | 2452 | + |

| | | | | | |
|---|---|---|---|---|---|
| VN1 | V1R23 | pri299568_MYRFA_USYD1HAP299568 | 21148 | 22071 | + |
| VN1 | V1R24 | pri300030_MYRFA_USYD1HAP300030 | 1128 | 244 | - |
| VN1 | V1R25 | pri300468_MYRFA_USYD1HAP300468 | 4535 | 3726 | - |
| VN1 | V1R26 | pri301800_MYRFA_USYD1HAP301800 | 1642 | 2628 | + |
| VN1 | V1R27 | pri301975_MYRFA_USYD1HAP301975 | 4921 | 5457 | + |
| VN1 | V1R28 | pri301975_MYRFA_USYD1HAP301975 | 15861 | 16853 | + |
| VN1 | V1R29 | pri301975_MYRFA_USYD1HAP301975 | 35450 | 36433 | + |
| VN1 | V1R110-partial | pri302109_MYRFA_USYD1HAP302109 | 34964 | 34137 | - |
| VN1 | V1R30 | pri302132_MYRFA_USYD1HAP302132 | 31322 | 30438 | - |
| VN1 | V1R31 | pri302132_MYRFA_USYD1HAP302132 | 58935 | 57991 | - |
| VN1 | V1R32 | pri303261_MYRFA_USYD1HAP303261 | 30741 | 31541 | + |
| VN1 | V1R111-partial | pri304051_MYRFA_USYD1HAP304051 | 12 | 449 | + |
| VN1 | V1R33 | pri304075_MYRFA_USYD1HAP304075 | 6023 | 6973 | + |
| VN1 | V1R34 | pri304075_MYRFA_USYD1HAP304075 | 24325 | 23441 | - |
| VN1 | V1R35 | pri304347_MYRFA_USYD1HAP304347 | 16797 | 15838 | - |
| VN1 | V1R36 | pri304347_MYRFA_USYD1HAP304347 | 35522 | 34602 | - |
| VN1 | V1R37 | pri304347_MYRFA_USYD1HAP304347 | 53386 | 52382 | - |
| VN1 | V1R38 | pri304481_MYRFA_USYD1HAP304481 | 5529 | 4558 | - |
| VN1 | V1R39 | pri304481_MYRFA_USYD1HAP304481 | 15141 | 14278 | - |
| VN1 | V1R40 | pri304813_MYRFA_USYD1HAP304813 | 31525 | 32502 | + |
| VN1 | V1R41 | pri304813_MYRFA_USYD1HAP304813 | 39561 | 40460 | + |
| VN1 | V1R42 | pri304813_MYRFA_USYD1HAP304813 | 59980 | 60942 | + |
| VN1 | V1R43 | pri305443_MYRFA_USYD1HAP305443 | 27904 | 27023 | - |
| VN1 | V1R44 | pri306318_MYRFA_USYD1HAP306318 | 10963 | 11859 | + |
| VN1 | V1R45 | pri306318_MYRFA_USYD1HAP306318 | 32398 | 33282 | + |
| VN1 | V1R46 | pri306318_MYRFA_USYD1HAP306318 | 54912 | 55862 | + |
| VN1 | V1R47 | pri306318_MYRFA_USYD1HAP306318 | 99392 | 98508 | - |
| VN1 | V1R48 | pri3067_MYRFA_USYD2HAP3067 | 34559 | 33660 | - |
| VN1 | V1R49 | pri3067_MYRFA_USYD2HAP3067 | 59028 | 59915 | + |
| VN1 | V1R50 | pri306740_MYRFA_USYD1HAP306740 | 49562 | 48738 | - |
| VN1 | V1R51 | pri306740_MYRFA_USYD1HAP306740 | 76546 | 75722 | - |
| VN1 | V1R52 | pri306837_MYRFA_USYD1HAP306837 | 12314 | 13288 | + |
| VN1 | V1R53 | pri306917_MYRFA_USYD1HAP306917 | 564 | 1472 | + |
| VN1 | V1R54 | pri307258_MYRFA_USYD1HAP307258 | 5699 | 4713 | - |
| VN1 | V1R55 | pri307418_MYRFA_USYD1HAP307418 | 16978 | 16085 | - |
| VN1 | V1R56 | pri307418_MYRFA_USYD1HAP307418 | 35778 | 34870 | - |
| VN1 | V1R57 | pri307418_MYRFA_USYD1HAP307418 | 48305 | 47394 | - |
| VN1 | V1R58 | pri307841_MYRFA_USYD1HAP307841 | 14297 | 13386 | - |
| VN1 | V1R59 | pri307968_MYRFA_USYD1HAP307968 | 18077 | 19027 | + |
| VN1 | V1R60 | pri307968_MYRFA_USYD1HAP307968 | 48232 | 49206 | + |
| VN1 | V1R61 | pri308403_MYRFA_USYD1HAP308403 | 6604 | 5720 | - |
| VN1 | V1R62 | pri308445_MYRFA_USYD1HAP308445 | 17656 | 18564 | + |
| VN1 | V1R63 | pri308459_MYRFA_USYD1HAP308459 | 8581 | 9567 | + |
| VN1 | V1R64 | pri308459_MYRFA_USYD1HAP308459 | 25916 | 26893 | + |
| VN1 | V1R65 | pri309576_MYRFA_USYD1HAP309576 | 11572 | 12525 | + |
| VN1 | V1R66 | pri309576_MYRFA_USYD1HAP309576 | 34894 | 35880 | + |
| VN1 | V1R67 | pri309576_MYRFA_USYD1HAP309576 | 63376 | 64356 | + |
| VN1 | V1R68 | pri309576_MYRFA_USYD1HAP309576 | 85012 | 85983 | + |
| VN1 | V1R69 | pri309576_MYRFA_USYD1HAP309576 | 102660 | 101737 | - |
| VN1 | V1R70 | pri309908_MYRFA_USYD1HAP309908 | 1611 | 715 | - |

| | | | | | |
|---|---|---|---|---|---|
| VN1 | V1R71 | pri310793_MYRFA_USYD1HAP310793 | 5476 | 4616 | - |
| VN1 | V1R72 | pri310793_MYRFA_USYD1HAP310793 | 5503 | 4613 | - |
| VN1 | V1R73 | pri310793_MYRFA_USYD1HAP310793 | 21494 | 20625 | - |
| VN1 | V1R74 | pri310793_MYRFA_USYD1HAP310793 | 36095 | 35229 | - |
| VN1 | V1R75 | pri3501_MYRFA_USYD1HAP3501 | 241028 | 240171 | - |
| VN1 | V1R76 | pri3699_MYRFA_USYD2HAP3699 | 89738 | 88914 | - |
| VN1 | V1R77 | pri3699_MYRFA_USYD2HAP3699 | 166962 | 166051 | - |
| VN1 | V1R78 | pri3870_MYRFA_USYD2HAP3870 | 267820 | 268767 | + |
| VN1 | V1R79 | pri4445_MYRFA_USYD1HAP4445 | 42386 | 41535 | - |
| VN1 | V1R80 | pri4445_MYRFA_USYD1HAP4445 | 52873 | 51980 | - |
| VN1 | V1R81 | pri4445_MYRFA_USYD1HAP4445 | 67718 | 66834 | - |
| VN1 | VR182 | pri4445_MYRFA_USYD1HAP4445 | 99574 | 98678 | - |
| VN1 | V1R83 | pri4687_MYRFA_USYD2HAP4687 | 106015 | 106935 | + |
| VN1 | V1R84 | pri5165_MYRFA_USYD1HAP5165 | 23677 | 22793 | - |
| VN1 | V1R85 | pri5165_MYRFA_USYD1HAP5165 | 41472 | 40495 | - |
| VN1 | V1R86 | pri5165_MYRFA_USYD1HAP5165 | 75030 | 74107 | - |
| VN1 | V1R87 | pri52_MYRFA_USYD1HAP52 | 7657 | 8577 | + |
| VN1 | V1R88 | pri549564_MYRFA_USYD1HAP549564 | 9483 | 8581 | - |
| VN1 | VAR112-partial | pri552459_MYRFA_USYD1HAP552459 | 1 | 681 | + |
| VN1 | V1R89 | pri575070_MYRFA_USYD1HAP575070 | 103 | 972 | + |
| VN1 | V1R90 | pri575070_MYRFA_USYD1HAP575070 | 17439 | 16549 | - |
| VN1 | VR191 | pri575071_MYRFA_USYD1HAP575071 | 17422 | 16592 | - |
| VN1 | VR192 | pri575078_MYRFA_USYD1HAP575078 | 20917 | 20000 | - |
| VN1 | V1R93 | pri575091_MYRFA_USYD1HAP575091 | 1198 | 2064 | + |
| VN1 | V1R94 | pri5980_MYRFA_USYD2HAP5980 | 261966 | 261076 | - |
| VN1 | V1R95 | pri631_MYRFA_USYD2HAP631 | 178211 | 177390 | - |
| VN1 | V1R96 | pri631_MYRFA_USYD2HAP631 | 189992 | 189186 | - |
| VN1 | V1R97 | pri631_MYRFA_USYD2HAP631 | 190058 | 189180 | - |
| VN1 | V1R98 | pri631_MYRFA_USYD2HAP631 | 213919 | 213020 | - |
| VN1 | V1R99 | pri631_MYRFA_USYD2HAP631 | 230579 | 229686 | - |
| VN1 | V1R100 | pri7389_MYRFA_USYD1HAP7389 | 20525 | 19638 | - |
| VN1 | V1R101 | pri7389_MYRFA_USYD1HAP7389 | 31362 | 30574 | - |
| VN1 | V1R102 | pri7389_MYRFA_USYD1HAP7389 | 31434 | 30574 | - |
| VN1 | V1R103 | pri7389_MYRFA_USYD1HAP7389 | 50404 | 49544 | - |
| VN1 | V1R104 | pri7389_MYRFA_USYD1HAP7389 | 60166 | 59315 | - |
| VN1 | V1R105 | pri7389_MYRFA_USYD1HAP7389 | 78621 | 77761 | - |
| VN1 | V1R106 | pri7389_MYRFA_USYD1HAP7389 | 87083 | 86223 | - |
| VN1 | V1R107 | pri7389_MYRFA_USYD1HAP7389 | 105463 | 106335 | + |
| VN1 | V1R108 | pri7643_MYRFA_USYD2HAP7643 | 2974 | 3963 | + |
| VN1 | V1R109 | pri8130_MYRFA_USYD1HAP8130 | 6573 | 7526 | + |
| VN1 | V1R1p | pri198488_MYRFA_USYD1HAP198488 | 981 | 325 | - |
| VN1 | V1R1p | pri217912_MYRFA_USYD1HAP217912 | 271 | 26 | - |
| VN1 | V1R1p | pri242107_MYRFA_USYD1HAP242107 | 1377 | 2120 | + |
| VN1 | V1R1p | pri261177_MYRFA_USYD1HAP261177 | 1179 | 1772 | + |
| VN1 | V1R1p | pri268596_MYRFA_USYD1HAP268596 | 51 | 908 | + |
| VN1 | V1R1p | pri270150_MYRFA_USYD1HAP270150 | 1537 | 2385 | + |
| VN1 | V1R1p | pri271592_MYRFA_USYD1HAP271592 | 6560 | 5940 | - |
| VN1 | V1R1p | pri275593_MYRFA_USYD1HAP275593 | 1650 | 1276 | - |
| VN1 | V1R1p | pri276995_MYRFA_USYD1HAP276995 | 1960 | 1063 | - |
| VN1 | V1R1p | pri282164_MYRFA_USYD1HAP282164 | 1531 | 1052 | - |

| | | | | | |
|---|---|---|---|---|---|
| VN1 | V1R1p | pri284998_MYRFA_USYD1HAP284998 | 4653 | 5150 | + |
| VN1 | V1R1p | pri287032_MYRFA_USYD1HAP287032 | 8189 | 7470 | - |
| VN1 | V1R1p | pri289009_MYRFA_USYD1HAP289009 | 1109 | 1330 | + |
| VN1 | V1R1p | pri289642_MYRFA_USYD1HAP289642 | 659 | 3 | - |
| VN1 | V1R1p | pri295589_MYRFA_USYD1HAP295589 | 6852 | 6046 | - |
| VN1 | V1R1p | pri295589_MYRFA_USYD1HAP295589 | 20809 | 19919 | - |
| VN1 | V1R1p | pri295589_MYRFA_USYD1HAP295589 | 42047 | 41131 | - |
| VN1 | V1R1p | pri296644_MYRFA_USYD1HAP296644 | 17815 | 17563 | - |
| VN1 | V1R1p | pri300462_MYRFA_USYD1HAP300462 | 6754 | 5828 | - |
| VN1 | V1R1p | pri302109_MYRFA_USYD1HAP302109 | 18093 | 17231 | - |
| VN1 | V1R1p | pri302109_MYRFA_USYD1HAP302109 | 46637 | 46074 | - |
| VN1 | V1R1p | pri303151_MYRFA_USYD1HAP303151 | 800 | 72 | - |
| VN1 | V1R1p | pri303151_MYRFA_USYD1HAP303151 | 19785 | 19022 | - |
| VN1 | V1R1p | pri303261_MYRFA_USYD1HAP303261 | 53393 | 53839 | + |
| VN1 | V1R1p | pri304481_MYRFA_USYD1HAP304481 | 18508 | 17978 | - |
| VN1 | V1R1p | pri304813_MYRFA_USYD1HAP304813 | 3636 | 4610 | + |
| VN1 | V1R1p | pri306318_MYRFA_USYD1HAP306318 | 90827 | 89877 | - |
| VN1 | V1R1p | pri3067_MYRFA_USYD2HAP3067 | 59169 | 59978 | - |
| VN1 | V1R1p | pri308445_MYRFA_USYD1HAP308445 | 36391 | 35456 | - |
| VN1 | V1R1p | pri309401_MYRFA_USYD1HAP309401 | 2048 | 1506 | - |
| VN1 | V1R1p | pri3699_MYRFA_USYD2HAP3699 | 123751 | 122882 | - |
| VN1 | V1R1p | pri3699_MYRFA_USYD2HAP3699 | 136657 | 136178 | - |
| VN1 | V1R1p | pri3711_MYRFA_USYD1HAP3711 | 4242 | 4955 | + |
| VN1 | V1R1p | pri3870_MYRFA_USYD2HAP3870 | 277579 | 278391 | + |
| VN1 | V1R1p | pri4187_MYRFA_USYD2HAP4187 | 3629 | 2773 | - |
| VN1 | V1R1p | pri4445_MYRFA_USYD1HAP4445 | 14056 | 13157 | - |
| VN1 | V1R1p | pri4445_MYRFA_USYD1HAP4445 | 27837 | 27088 | - |
| VN1 | V1R1p | pri4445_MYRFA_USYD1HAP4445 | 33358 | 32468 | - |
| VN1 | V1R1p | pri4445_MYRFA_USYD1HAP4445 | 109602 | 108991 | - |
| VN1 | V1R1p | pri5165_MYRFA_USYD1HAP5165 | 2040 | 1474 | - |
| VN1 | V1R1p | pri5165_MYRFA_USYD1HAP5165 | 16023 | 15340 | - |
| VN1 | V1R1p | pri552461_MYRFA_USYD1HAP552461 | 2192 | 2875 | + |
| VN1 | V1R1p | pri575069_MYRFA_USYD1HAP575069 | 8560 | 7693 | - |
| VN1 | V1R1p | pri575069_MYRFA_USYD1HAP575069 | 21034 | 20118 | - |
| VN1 | V1R1p | pri575078_MYRFA_USYD1HAP575078 | 8448 | 7581 | - |
| VN1 | V1R1p | pri631_MYRFA_USYD2HAP631 | 156830 | 155931 | - |
| VN1 | V1R1p | pri631_MYRFA_USYD2HAP631 | 167601 | 166945 | - |
| VN1 | V1R1p | pri631_MYRFA_USYD2HAP631 | 203117 | 202386 | - |
| VN1 | V1R1p | pri631_MYRFA_USYD2HAP631 | 243216 | 242425 | - |
| VN1 | V1R1p | pri7697_MYRFA_USYD1HAP7697 | 12118 | 11153 | - |
| VN2 | V2R_partial | pri1901_MYRFA_USYD1HAP1901 | 287143 | 318774 | + |
| VN2 | V2R | pri2104_MYRFA_USYD2HAP2104 | 308374 | 324870 | + |
| VN2 | V2R | pri2104_MYRFA_USYD2HAP2104 | 336559 | 360548 | + |
| VN2 | V2R_partial | pri2104_MYRFA_USYD2HAP2104 | 367990 | 380432 | + |
| VN2 | V2R_partial | pri2104_MYRFA_USYD2HAP2104 | 394995 | 401976 | + |
| VN2 | V2R | pri2104_MYRFA_USYD2HAP2104 | 415107 | 439708 | + |
| VN2 | V2R_partial | pri2547_MYRFA_USYD1HAP2547 | 376041 | 369800 | - |
| VN2 | V2R_partial | pri255135_MYRFA_USYD1HAP255135 | 13965 | 8828 | - |
| VN2 | V2R | pri266717_MYRFA_USYD1HAP266717 | 12852 | 5702 | - |
| VN2 | V2R | pri287859_MYRFA_USYD1HAP287859 | 22569 | 3748 | - |

| | | | | | |
|---|---|---|---|---|---|
| VN2 | V2R_partial | pri291047_MYRFA_USYD1HAP291047 | 2412 | 3 | - |
| VN2 | V2R_partial | pri300513_MYRFA_USYD1HAP300513 | 14143 | 9153 | - |
| VN2 | V2R_partial | pri301297_MYRFA_USYD1HAP301297 | 4107 | 13472 | + |
| VN2 | V2R | pri302618_MYRFA_USYD1HAP302618 | 27594 | 5217 | - |
| VN2 | V2R_partial | pri306650_MYRFA_USYD1HAP306650 | 1057 | 8008 | + |
| VN2 | V2R_partial | pri306650_MYRFA_USYD1HAP306650 | 32970 | 42824 | + |
| VN2 | V2R_partial | pri308657_MYRFA_USYD1HAP308657 | 71350 | 59671 | - |
| VN2 | V2R_partial | pri3196_MYRFA_USYD2HAP3196 | 10668 | 1746 | - |
| VN2 | V2R_partial | pri4553_MYRFA_USYD2HAP4553 | 183458 | 182547 | - |
| VN2 | V2R_partial | pri482648_MYRFA_USYD1HAP482648 | 802 | 3136 | + |
| VN2 | V2R | pri5071_MYRFA_USYD2HAP5071 | 202985 | 231060 | + |
| VN2 | V2R | pri5071_MYRFA_USYD2HAP5071 | 313434 | 289406 | - |
| VN2 | V2Rp | pri2104_MYRFA_USYD2HAP2104 | 410723 | 457611 | + |
| VN2 | V2Rp | pri287859_MYRFA_USYD1HAP287859 | 47913 | 29585 | - |
| VN2 | V2Rp | pri292225_MYRFA_USYD1HAP292225 | 12820 | 2811 | - |
| VN2 | V2Rp | pri5071_MYRFA_USYD2HAP5071 | 285614 | 182873 | - |
| VN2 | V2Rp | pri6986_MYRFA_USYD2HAP6986 | 9533 | 24741 | |
| Aquaporin | AQP2 | pri284316_MYRFA_USYD1HAP284316 | 12063 | 22237 | + |
| Aquaporin | AQP5 | pri284316_MYRFA_USYD1HAP284316 | 32393 | 35334 | + |
| Aquaporin | AQP6 | pri300950_MYRFA_USYD1HAP300950 | 93854 | 90539 | - |
| Aquaporin | AQP4 | pri302576_MYRFA_USYD1HAP302576 | 11007 | 24840 | - |
| Aquaporin | AQP12-partial | pri3559_MYRFA_USYD2HAP3559 | 25424 | 28493 | - |
| Aquaporin | AQP3 | pri4880_MYRFA_USYD1HAP4880 | 50362 | 59029 | - |
| Aquaporin | AQP7 | pri5086_MYRFA_USYD1HAP5086 | 14717 | 48966 | - |
| Aquaporin | AQP10 | pri5319_MYRFA_USYD1HAP5319 | 49660 | 54287 | + |
| Aquaporin | AQP1 | pri563438_MYRFA_USYD1HAP563438 | 12080 | 6880 | - |
| Aquaporin | AQP11 | pri5767_MYRFA_USYD2HAP5767 | 25746 | 43538 | + |
| Aquaporin | AQP8 | pri6070_MYRFA_USYD1HAP6070 | 132170 | 126193 | - |
| Aquaporin | AQP9 | pri6897_MYRFA_USYD2HAP6897 | 3702 | 65635 | - |
